# Cell type-independent timekeeping gene modules enable embryonic stage prediction in zebrafish

**DOI:** 10.1101/2025.11.12.688039

**Authors:** Rupa Kanchi, Sandra L Grimm, Divya Vella, Richard Saoud, Tanmay Gandhi, Amrit Koirala, Ailen Cervino, Jacalyn MacGowan, Cristian Coarfa, Margot Kossmann Williams

**Affiliations:** Molecular and Cellular Biology Department, Baylor College of Medicine, Houston; Dan L Duncan Comprehensive Cancer Center, Baylor College of Medicine, Houston; Center for Precision Environmental Health, Baylor College of Medicine, Houston; Molecular Virology and Microbiology Department, Baylor College of Medicine, Houston

## Abstract

Gene expression changes across embryonic development reflect both differentiation and genes whose expression varies strictly with developmental time, independent of cell type. Multiple embryonic timing systems set the onset and pace of developmental events, and blocking transcription arrests many of these events. However, the genes comprising the vertebrate embryonic timing system(s) remain largely unknown. To identify genes whose expression changes with time alone, we examine naive zebrafish embryonic explants that form only two tissue types yet maintain developmental timing, thus uncoupling developmental age from most differentiation programs. By comparing longitudinal gene expression in naïve explants with Nodal-induced explants that differentiate into all three germ layers, we identify “timekeeping” genes whose temporal expression patterns vary strictly with developmental age. Consensus clustering of temporally dynamic genes identified 20 gene clusters, termed “chrono-constitutive modules” (**CCMs**), that maintain distinct schedules of expression regardless of tissue type. These CCM trajectories are similar in intact zebrafish embryos and single embryonic cells of multiple distinct lineages. Enrichment analysis of microRNA targets and transcription factor regulons within the CCMs further reveal distinct putative regulators of several modules. Strikingly, CCM expression patterns are also largely conserved during early development of another fish species, Japanese medaka. Machine learning models trained on only zebrafish CCM transcript levels accurately predict the developmental age of embryonic explants, intact embryos, and even individual embryonic cells, demonstrating their utility in developmental timekeeping. These results support the existence of transcriptional timekeeping during early development and demonstrate its utility in embryonic stage prediction.

## Introduction

Embryonic development is remarkably reproducible, exhibiting great precision in the timing of cell divisions, gene expression, cell type specification, and tissue morphogenesis between individuals of a species, with very little variation in the onset and pace of many developmental events. This remarkable ability of embryos to keep time is attributed to multiple developmental timing mechanisms, often referred to as embryonic “clocks” or “timers” (*1, 2*). Evidence suggests that many such mechanisms function during early development. For example, early embryonic cleavages are under the control of the Cyclin “clock” (*3*). The resulting increase in nuclear/cytoplasmic ratio (*4, 5*) (*6-9*) then activates the mid-blastula transition (MBT) – characterized by zygotic genome activation, slowed and asynchronous cell divisions, and cell motility onset. The onset of gastrulation cell movements, germ layer specification, and differentiation are also precisely timed in anamniote species, but these events are uncoupled from cell number and tissue volume (*10-19*), necessitating yet another timing mechanism. Notably, each of these timing systems functions independently of one another. Cell divisions *per se* are not required for MBT or differentiation (*4, 10-14, 20*), for example, and the time of MBT does not determine the time of gastrulation onset (*15, 19, 21-23*). This demonstrates the existence of multiple timing systems in even the earliest embryos, each driven by distinct molecular mechanisms. While the molecular basis for some developmental timers (like those driving MBT) has been at least partially elucidated, others (like gastrulation onset) remain largely unknown. The molecular basis of embryonic timing systems remains a fundamental open question in developmental biology.

One input into – and output of - many embryonic timing systems is transcription. New gene expression is required for the advancement or progression of some timers, such as those controlling gastrulation onset and differentiation (*4, 10, 12, 24*). Activation of transcription is also the predominant indicator for zygotic genome activation, but gene expression continues to change dramatically throughout embryogenesis, marking developmental progression. One obvious reason for this is that, as an embryo develops, its constituent cells differentiate into an increasingly diverse array of cell and tissue types each expressing a complement of tissue-specific genes. However, we hypothesize that temporal changes also reflect a second class of strictly “timekeeping” genes whose expression varies only with innate developmental time, independent of cell or tissue type. How many genes fall into each of these categories is unclear, and identifying a suite of embryonic timekeeping genes is the first step to understanding the transcriptional inputs and outputs of poorly defined developmental timing systems.

Both differentiation and innate timekeeping genes must be reflected in bulk transcriptomic profiling of embryos. Developmental time-courses of gene expression have been reported for mouse (*25, 26*), chick (*27, 28*), *Xenopus tropicalis* and *laevis* (*29, 30*), medaka (*31*), *Fundulus* (*32*), zebrafish (*33-37*), *Drosophila* (*38-40*), *C. elegans* (*41-43*), and even humans (*44-46*). In these whole embryos, transcripts can be clustered according to their temporal expression profiles, reflecting different expression (and/or degradation) dynamics of groups of genes across development. Transcriptomic profiles are unique to each developmental stage within a given species and contain sufficient timing information to infer age/stage, demonstrated by several bioinformatic approaches used to stage or order embryos based on bulk transcriptomes alone (*37, 38, 47-49*). Notably, differentiation signatures provided the majority of staging information for one such tool, rather than strictly timekeeping genes (*49*). Stage/age predictors (or refiners) were also built from single-cell transcriptomes (*50-53*), some of which used ribosome or cell cycle genes to infer a “general transcriptomic clock” (*51, 52*) that was not based on differentiation of specific cell types. Indeed, consideration of both new gene expression and degradation of maternal transcripts in single cells of zebrafish embryos revealed that mRNA dynamics are constant between distinct cell lineages for the vast majority of genes (*54*). This suggests that most temporal expression dynamics are cell type-agnostic, at least during pre-gastrulation stages, which together with transcriptomic staging tools highlights the feasibility of temporally dynamic genes to convey developmental timing information.

We sought to characterize a core set of embryonic timekeeping genes whose expression varies strictly with developmental progression in a cell type-agnostic way. To this end, we employed zebrafish embryonic explants in which “timekeeping” gene expression is maintained in the absence of most tissue types. When Nodal signaling is activated, these explants specify multiple tissues of all three germ layers (*55, 56*). Critically, they also maintain the precise timing of cell type specification and gastrulation morphogenesis observed in whole embryos (*55, 57*), demonstrating that their intrinsic developmental timer is intact. In the absence of exogenous signals, these explants make only two tissue types, non-neural ectoderm and enveloping layer (*58*). Therefore, genes with similar expression dynamics in both activated and naïve explants are likely to represent cell type-agnostic timekeeping genes and exclude differentiation signatures of the many cell types present in Nodal-activated explants and intact embryos.

By comparing longitudinal transcriptional profiles of Nodal-activated and naïve zebrafish explants from blastula to neurula stages (*59*), we identified 24 distinct clusters of temporal expression patterns, or “chrono-modules”. Of these, 20 were nearly identical between naïve and Nodal-activated explants, suggesting that most developmental genes are “chrono-constitutive” and maintain their schedule of expression regardless of cell type. Expression of these 20 chrono-constitutive modules (**CCMs**) was highly similar between zebrafish explants, intact embryos, and single embryonic cells of the same stages, validating their significance to *in vivo* development. Many modules associated with specific signaling pathways, transcription factors, and microRNA families, hinting at distinct molecular regulation of each. Expression trajectories of these CCMs were also largely conserved in another fish species, Japanese medaka. Finally, we found that machine learning models trained on CCMs alone accurately predicted the stage of explants, intact embryos, and even single cells, indicating they are sufficient to convey developmental timing information. Together, these data reveal the existence of a core set of evolutionarily conserved timekeeping genes during early development whose expression reflects developmental progression independent of differentiation.

## Results

### Zebrafish explants exhibit strictly stage-dependent gene expression

To identify genes whose temporal expression patterns change strictly with developmental time rather than cell type specification and differentiation, we examined our previously published gene expression profiling of naïve and Nodal-activated zebrafish explants at seven developmental stages (*59*)(**Fig. 1A**). These explants were generated by isolating and culturing the animal poles from 256- to 512-cell stage zebrafish embryos (*56, 60, 61*) injected with RNA encoding the Nodal ligand Cyclops (*ndr2*), a constitutively active Nodal receptor (*CA-acvr1b*\*), and uninjected controls. By 12 hours post fertilization (hpf), Nodal-induced explants express markers of all three germ layers and distinct populations of neuroectoderm and mesoderm (*55, 61*). By contrast, uninjected explants remain relatively naïve, expressing markers of non-neural ectoderm and enveloping layer but few markers of neuroectoderm, mesoderm, or endoderm (*55, 58, 60*). RNA was isolated from pooled explants of each condition when intact sibling embryos reached each of the following seven developmental stages: sphere (4 hpf), 30% epiboly (4.7 hpf), 50% epiboly (5.3 hpf), shield (6 hpf), 75% epiboly (8 hpf), 90% epiboly (9 hpf), and 2 somite (11 hpf) in biological triplicate (*59*) (**Fig. 1A**). Principal component analysis of all 63 transcriptomes showed that samples clustered predominantly with those of the same developmental stage, with little separation by their Nodal treatment status (**Fig. 1B**). This suggests that gene expression varies more with developmental stage than with development of specific cell types / lineages, supporting the notion of cell type-agnostic timekeeping gene expression.

**Figure 1.**
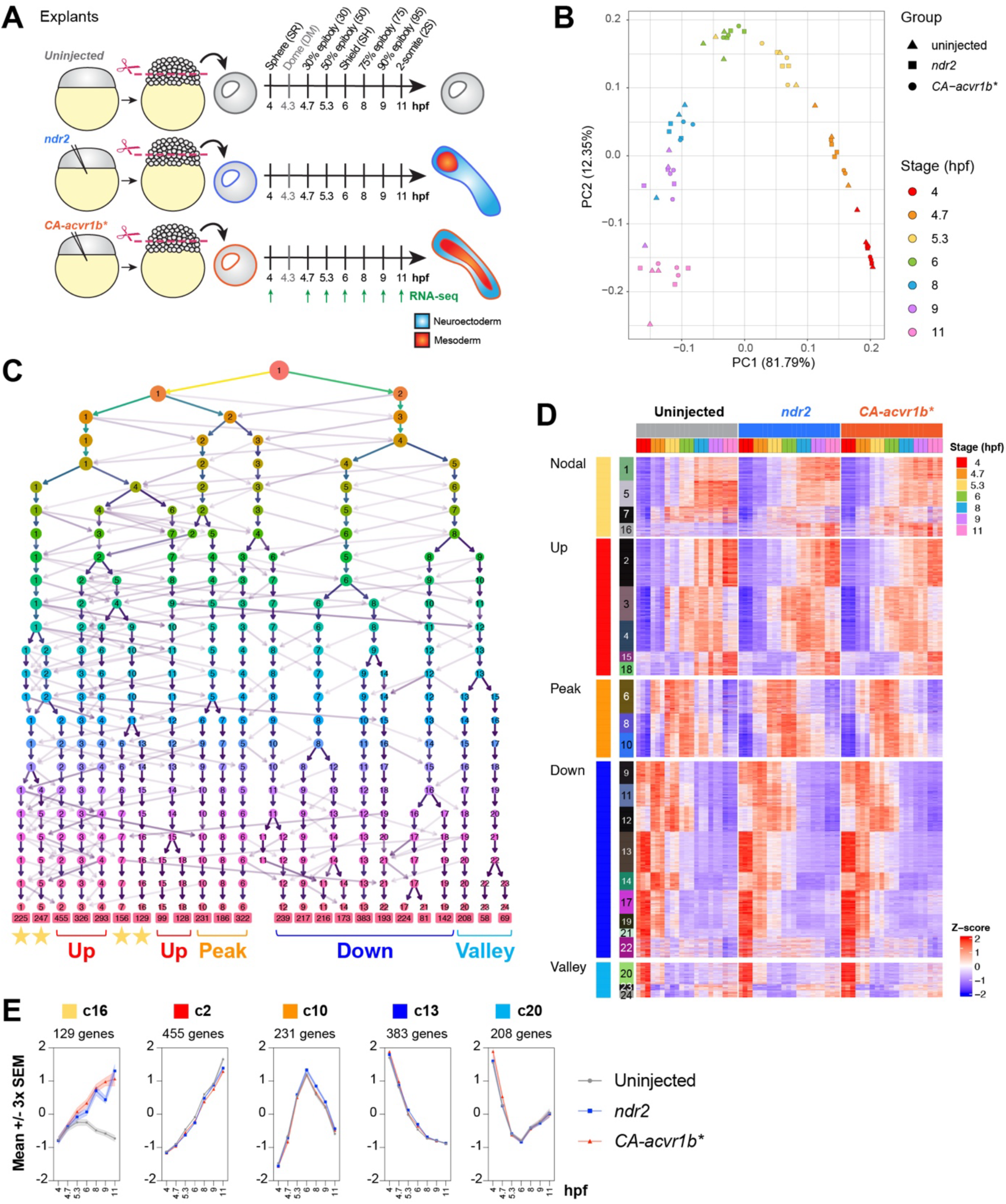
Consensus clustering of temporal gene expression profiles reveals chrono-constitutive gene modules. **A**) Schematic of longitudinal RNA-sequencing of zebrafish embryonic explants, including 7 time points and 3 biological replicates for each condition. **B**) Principal component analysis of the 63 explant RNA-seq samples. **C**) Clustering tree after applying consensus clustering, with number of modules varying from two to 24. Clusters with similar trajectory shapes (up, down, peak, valley) tend to be more closely related. **D**) Heatmap visualization of expression over time for the genes in each of 24 chrono-modules, grouped by trajectory shape. **E**) Summed z-score trajectories of five representative chrono-modules in uninjected, *ndr2*, and *CA-acvr1b*\* explant treatment groups. Diagram in (A) is adapted from (*55*). Based on longitudinal explant transcriptomic data published in (*59*).

**Supplementary Figure 1.**
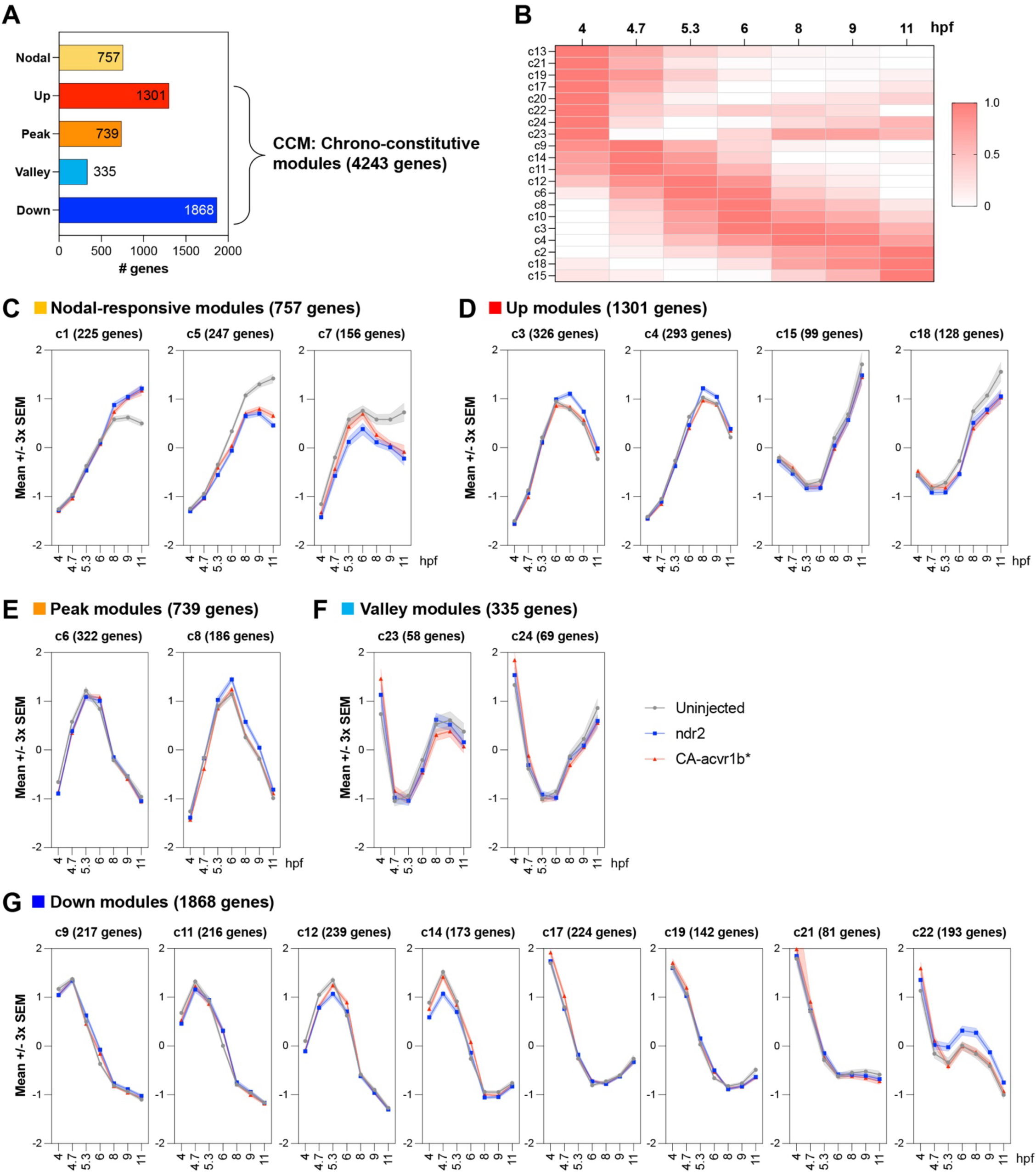
Temporal expression trajectories of chrono-modules. **A**) The number of (the top 5000 most temporally dynamic) genes within each of the five categories of chrono-modules. **B**) Scaled expression of the 20 chrono-constitutive modules ordered by timepoint of peak expression. **C-G**) Summed z-score trajectories of the indicated chrono-modules in uninjected, *ndr2*, and *CA-acvr1b*\* explant treatment groups.

### Clustering of temporal expression profiles reveals chrono-constitutive gene modules

To further examine temporal gene expression patterns within our zebrafish explants, we selected the 5000 most temporally variable genes and performed consensus clustering of temporal expression patterns across all explant conditions (**Fig. 1C**). Importantly, genes were clustered according to *changes* in their expression profiles over time (using a z-score transformation), regardless of *absolute* expression levels, resulting in 24 “chrono-modules” containing between 58 and 455 genes each. These modules clustered according to the general shapes of their expression trajectories over time, which we categorized as “up”, “down”, “peak”, and “valley”, with modules of the same general shape being more closely related on the clustering tree (**Fig. 1C-E, Supp Fig. 1A**). Most modules, even those categorized as “up” or “down”, contained peaks and/or valleys that differed in the stage at which their apex or nadir occurred and expression levels before and after the apex/nadir (**Fig. 1D-E, Supp Fig. 1B-G**).

Strikingly, the shapes of most gene module trajectories were essentially identical between the three explant conditions, with only four modules (out of 24) containing 757 genes (out of 5000) differing substantially between *ndr2*, *CA-acvr1b*\*, and uninjected explants (**Fig. 1D-E** and **Supp Fig. 1C**, “Nodal-responsive” clusters). Of these, modules 1 and 5 were similar between the two Nodal conditions which differed from uninjected explants, while modules 7 and 16 exhibited distinct temporal trajectories in each of the three explant conditions **Fig. 1D-E** and **Supp Fig. 1**). The most dramatic of these differences were in module 16, which resembled a small hill in uninjected, a jagged peak in *ndr2*, and a steep upward slope in *CA-acvr1b*\* explants (**Fig. 1E**). Module 16 contained many known Nodal-dependent genes (such as *tbxta, chrd, noto*, and *fgf8a*) and those involved in Nodal signaling itself (such as *ndr2* and *tdgf1*), highlighting this module as the predominant Nodal responsive cluster (**Supp Table 1**). The vast majority of genes included in our analysis (4243 of 5000), however, belonged to the remaining 20 modules which were highly similar between explant conditions (**Fig. 1D-E** and **Supp Fig. 1D-G**). We note that, although the shape of any given gene’s temporal expression trajectory (based on z-score transformation) is similar between conditions, absolute expression levels may not be. For example, *sox2* is within module 4 (a CCM) but we found that it is more strongly expressed in Nodal-induced than uninjected explants at the 2-somite stage (*55*)(**Supp Table 1**), although the shape of its expression trajectory is similar. *dlx3b*, a module 2 gene, is expressed at substantially higher levels in uninjected explants than in *ndr2* and *CA-acvr1b*\*, but increases monotonically in all conditions (**Supp Table 1**). This demonstrates that the temporal expression dynamics of genes within these clusters vary with developmental time, regardless of the explants’ constituent cell types. For this reason, we have termed these clusters “chrono-constitutive modules” (CCMs).

### Chrono-modules contain functionally distinct genes

To gain insight into the types of genes included within each chrono-module, we performed gene annotation enrichment analysis using Metascape (*62*) and found that many were enriched for specific biological processes (**Fig. 2, Supp Fig. 2, Supp Table 2**). As expected, Nodal-responsive modules were enriched for terms associated with Nodal-dependent embryonic processes like mesoderm and endoderm development, cell signaling, and morphogenesis (**Fig. 2A, Supp Fig. 2A**). Many CCMs, on the other hand, were enriched for terms associated with essential cellular functions like transcription, translation, and autophagy. For example, modules 3, 6, 8, 10, and 12, which all peak near the onset of gastrulation, were enriched for terms such as RNA processing and transcription (**Fig. 2B, Supp Fig. 2B-D**). Module 2, characterized by steadily increasing expression, was highly enriched for terms related to ribosome biogenesis and translation (**Fig. 2C**), and modules 20, 22, and 23 were enriched for genes involved in Ubiquitin-dependent protein degradation (**Fig. 2D-E**). However, some CCMs were enriched for non-housekeeping functions. Module 13, the only module to decrease monotonically throughout development, contained genes involved in blastoderm segmentation, P-body assembly, and negative regulation of translation (**Supp Fig. 2D**). This module includes maternally expressed genes like *buc*, *piwi2l*, and *zp3* that regulate oocyte and primordial germ cell development (**Supp Table 1**), suggesting that it represents maternal transcripts that are degraded over time. Additional examples include “central nervous system development” (module 15), “Rho GTPase cycle” (module 17), and “positive regulation of phosphatidylinositol 3-kinase signaling” (module 24) (**Supp Table 2**). Consistent with previous findings from intact embryos (*33*), these results indicate that genes with similar functions often exhibit coordinated temporal expression trajectories.

**Figure 2.**
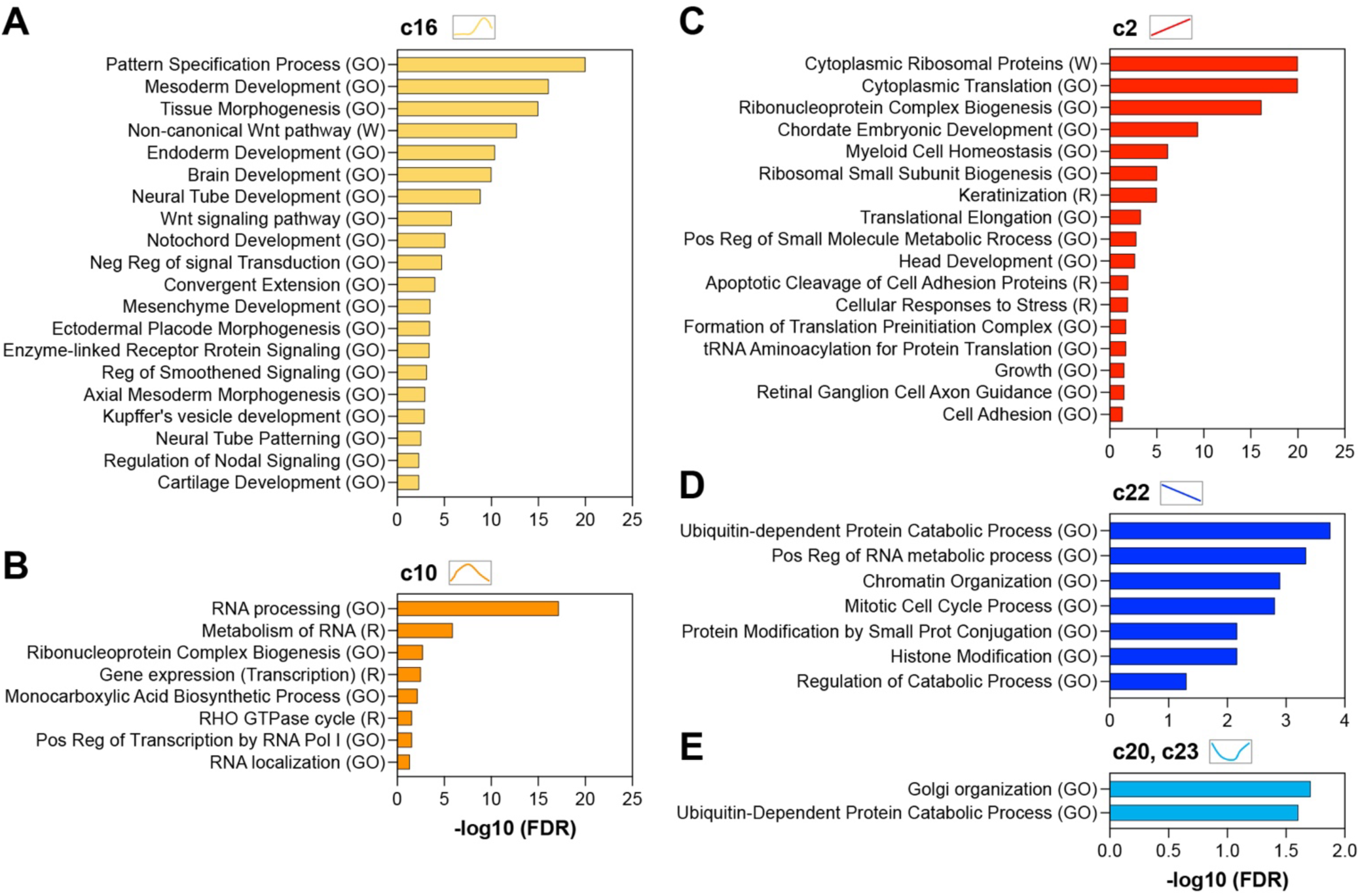
Functional enrichment of chrono-modules. Enriched gene annotation terms identified using Metascape are shown in select Nodal-responsive (A), peak (B), up (C), down (D), and valley (E) chrono-modules.

**Supplementary Figure 2.**
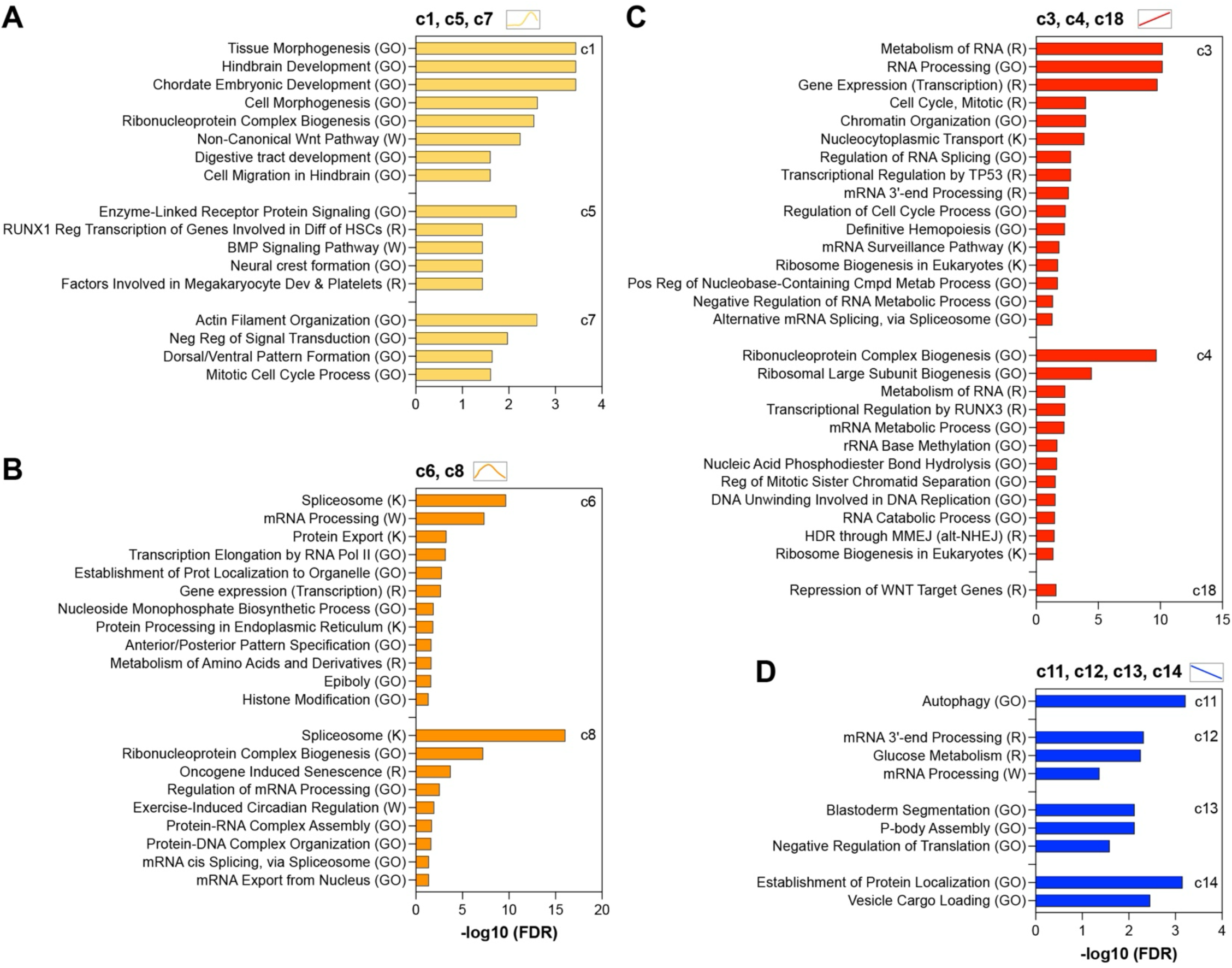
Functional enrichment of select chrono-modules. Enriched gene annotation terms identified using Metascape are shown in select Nodal-responsive (A), peak (B), up (C), and down (D), chrono-modules.

### Chrono-constitutive modules are concordant in intact zebrafish embryos

We next examined expression patterns of genes within all 24 chrono-modules across development in intact zebrafish embryos using a published bulk RNA-sequencing (RNA-seq) time-course (*33*). The stages included in this dataset were similar to those sampled in our explants, including 4.3, 5.3, 6, 8, and 11 hpf, and contained 4243 of the 5000 chrono-module genes selected from our explant dataset. When plotted across these timepoints, the shapes of most CCM gene modules were remarkably similar between our explants and whole embryos (**Fig. 3, Supp Fig. 3**, black graphs). For example, module 2 exhibited monotonic increases, module 13 decreased monotonically, and modules 20, 23, and 24 resembled valleys in all conditions. Notably, some groups of expression trajectories tended to match better between explants and embryos than others. As mentioned, valley-shaped CCMs and those whose expression decreased steadily at early stages were particularly consistent (**Fig. 3, Supp Fig. 3**, black graphs). However, many peak-shaped CCMs in explants were missing the pre-apex portion of the peak in intact embryos (see modules 6, 8, 9, 11, 12, and 14, **Supp Fig. 3C, E**, black graphs). Whether these differences reflect the slight disparity in stages profiled or real differences in gene expression is unclear. Notably, the authors of this whole embryo RNA-seq study also clustered genes according to their temporal expression patterns and identified gene modules that peaked at different stages (*33*). However, because the full dataset includes stages from 1-cell stage to 5 days post fertilization, these clusters and their shapes differ dramatically from those including only the subset of stages we selected for re-analysis.

**Figure 3.**
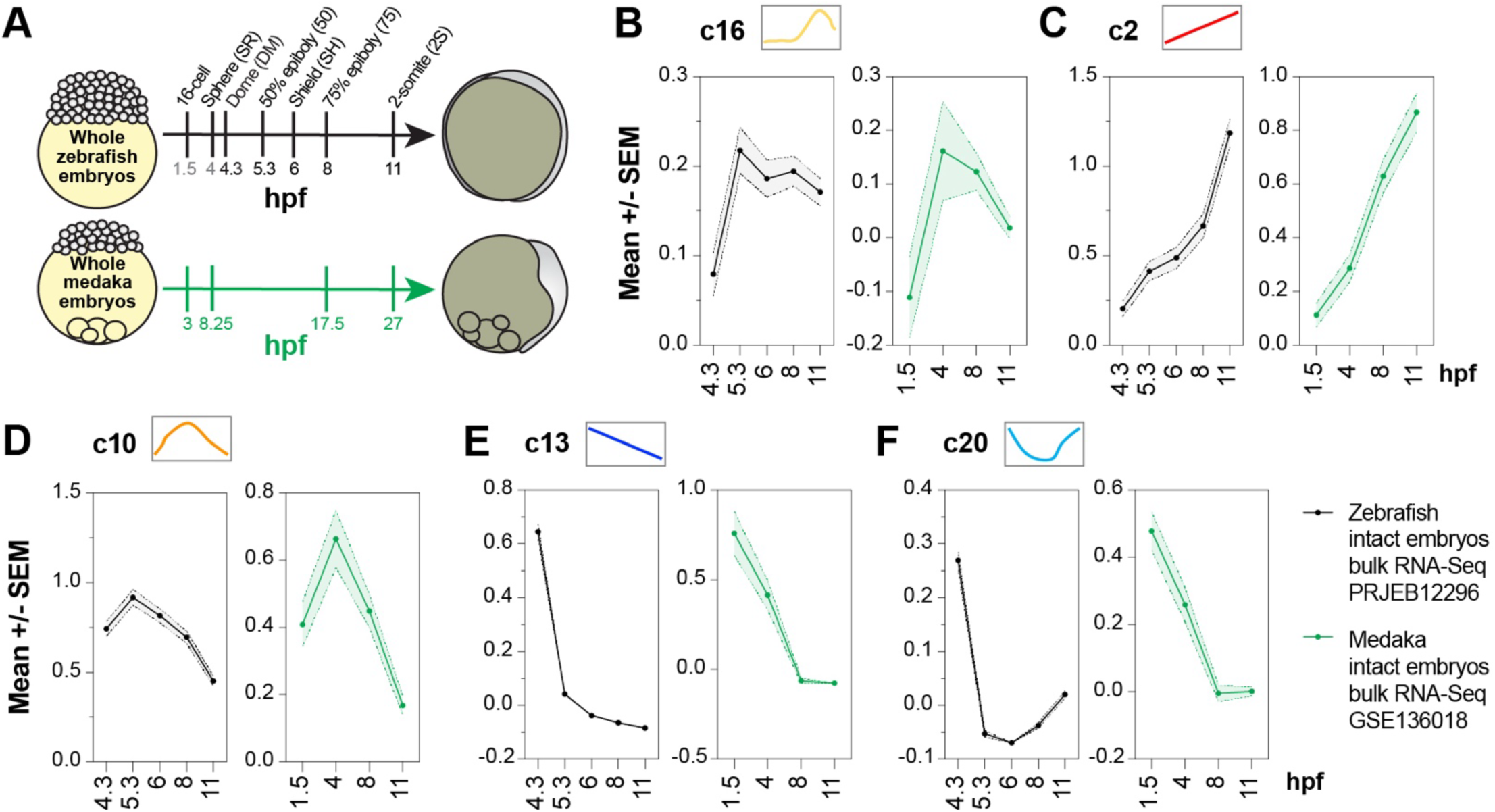
Chrono-modules are concordant in intact zebrafish embryos and conserved in medaka embryos. **A**) Schematic of developmental stages profiled in intact zebrafish and medaka embryos. **B-F**) Summed z-score trajectories of the indicated chrono-modules for intact zebrafish and medaka embryos. X axis shows hpf for zebrafish (black) or hpf of the nearest equivalent zebrafish stage for medaka (green). Based on published zebrafish and medaka embryo transcriptomes PRJEB12296 (*33*) and GSE136018 (*31*), respectively.

**Supplementary Figure 3.**
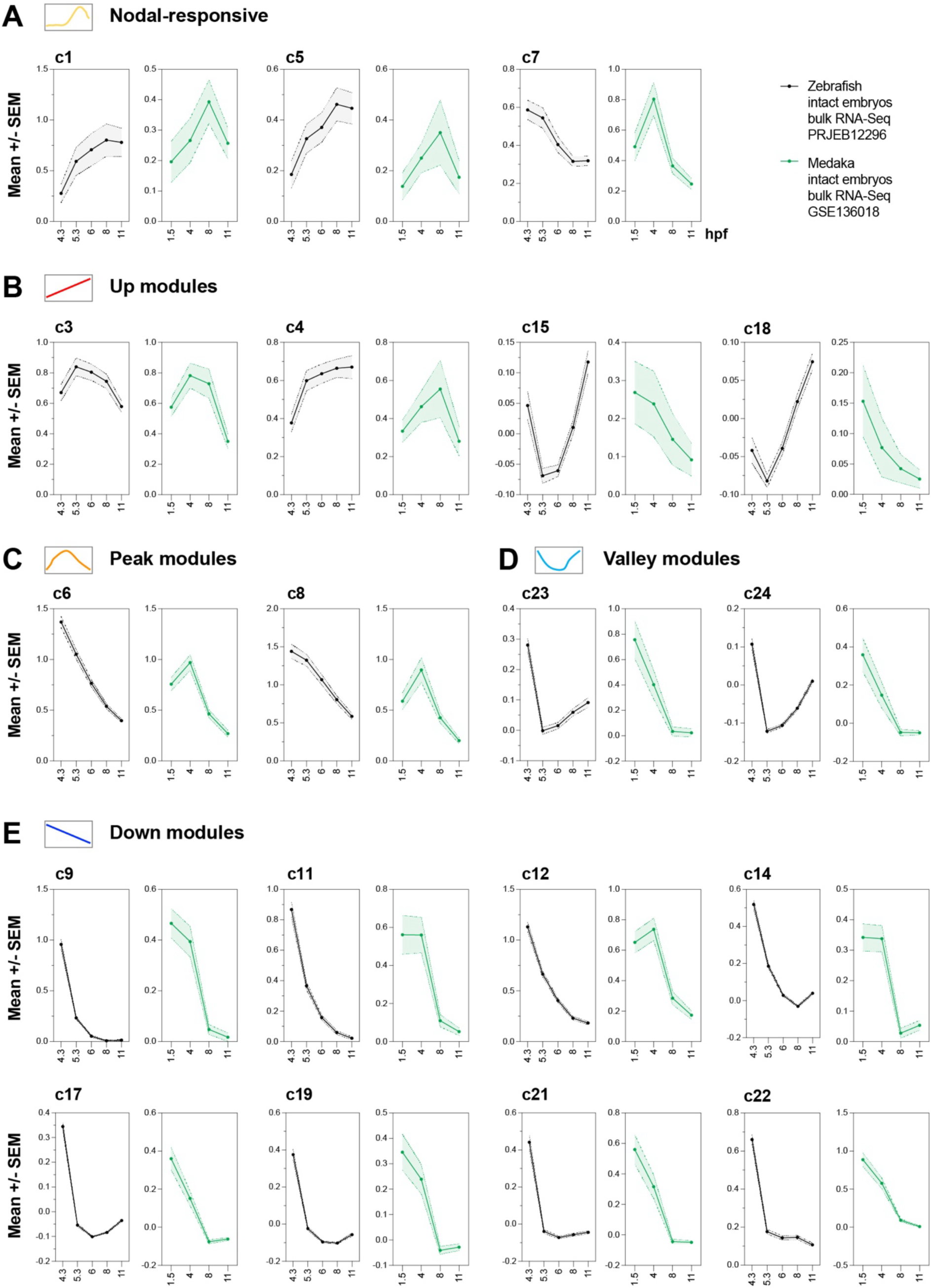
**Chrono-modules are concordant in intact zebrafish embryos and conserved in medaka embryos. A-E**) Summed z-score trajectories of the indicated chrono-modules for intact zebrafish (black) and medaka (green) embryos. X axis shows hpf for zebrafish or hpf of the nearest equivalent zebrafish stage for medaka. Based on published zebrafish and medaka embryo transcriptomes PRJEB12296 (*33*) and GSE136018 (*31*), respectively.

Unlike CCMs, temporal expression trajectories of Nodal-responsive gene modules were often quite different between explants and intact embryos. While module 5 in whole embryos more closely resembled Nodal-activated explants than uninjected, modules 1 and 16 in whole embryos looked much more like uninjected explants (**Fig. 3B, Supp Fig. 3A**, black graphs). This is somewhat unexpected, given that whole embryos exhibit Nodal signaling activity throughout blastula and gastrula stages (*63*). Module 7, on the other hand, resembled neither uninjected nor Nodal-induced explants. Together, these observations suggest that while chrono-constitutive gene expression within zebrafish embryos is generally well reflected in explants, Nodal-induced transcriptional changes are likely sensitive to the timing and method of Nodal activation. The similarity of strictly time-dependent gene expression trajectories between whole embryos and explants during these stages both validates our use of this *ex vivo* system for longitudinal profiling and highlights the consistency of CCM gene expression across developmental time.

### Chrono-constitutive gene expression is largely conserved in medaka embryos

Previous studies have observed similar temporal expression profiles of some developmental genes across even distantly related species (*25, 48, 64-66*). Indeed, the RAPToR tool can accurately stage embryos of one species using reference data from another (*49*). To test if our CCMs are conserved beyond zebrafish, we examined temporal expression patterns of homologs of these 5000 dynamic genes in embryos of another teleost fish, the Japanese medaka (*Oryzias latipes*). Using a publicly available longitudinal RNA-seq dataset encompassing four similar developmental stages (*31*), we identified medaka homologs for 3897 of the 5000 chrono-module genes and examined the expression of each homologous module over time. Although the trajectory shapes are coarser because the medaka dataset contains fewer timepoints, many modules were strikingly similar to those of zebrafish embryos and explants (**Fig. 3, Supp Fig. 3**, green graphs). Those with the most similarity between these species tended to be the “down” and “valley” modules (**Supp Fig. 3D-E**), whereas “up” modules diverged more, with two instead exhibiting decreasing expression (**Supp Fig. 3B**). Despite these few differences, these findings indicate that strictly time-dependent gene expression is largely conserved among closely related teleost species.

### Regulatory enrichment of chrono-modules

Our analysis demonstrates that CCM genes exhibit highly specific temporal expression trajectories, raising questions as to how their expression levels are regulated. Several mechanisms by which embryos control the level and/or time at which transcripts are expressed or degraded include transcription factor binding (*67, 68*), chromatin accessibility (*68-70*), promoter-enhancer associations (*71, 72*), transcriptional competence (*73*), intron length (*74, 75*), polymerase pausing (*76-78*), codon and 3’ UTR sequences (*54, 79*), and microRNA expression (*80, 81*), among others. To determine whether one or more of these mechanisms may explain expression patterns for any of our CCMs, we examined each module for enrichment of specific transcription factor binding sites (TFBSs) and predicted microRNA binding. Based on SwissRegulon annotations (*82*), we found that the constituent genes for twelve chrono-modules were enriched for at least one TFBS, and that each class of TF was often specific to a single module (**Fig. 4A, Supp Table 3**). Of the four Nodal-responsive modules, only module 16 showed significant enrichment of specific TFBSs. Consistent with its previous characterization as the Nodal-responsive cluster, module 16 was enriched for TFBSs for T-box transcription factors, many of which are themselves induced by Nodal (like Tbxta and Tbx16) (*83-85*) and have well described roles in mesoderm development (*86*). Among CCMs, module 12 was highly enriched for binding sites for Erf/Elk/Elf and Etv transcription factors, all belonging to the ETS domain-containing family of TFs.

**Figure 4.**
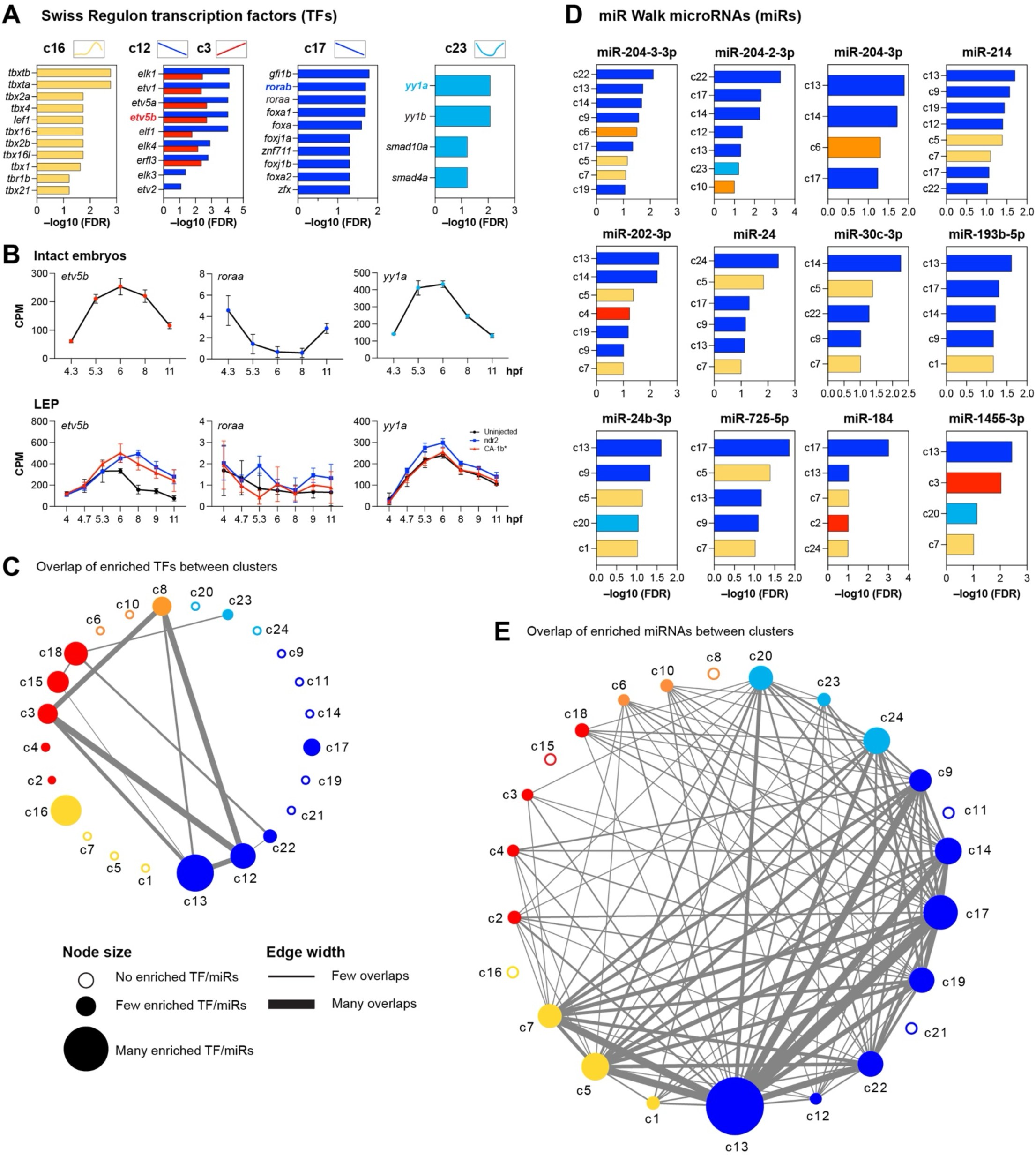
Chrono-modules are enriched for transcriptional and post-transcriptional regulators. **A**) Enrichment for transcription factor binding sites (TFBS) within the chrono-modules indicated, based on the SwissRegulon database. **B**) Temporal expression profiles of the transcription factors indicated in intact embryos (top) and all three explant conditions (bottom) mirror the profiles of chrono-modules (c3/12, c17, and c23) enriching for them. **C**) Network diagram of TFBS enrichment within chrono-modules. Nodes indicate the number of enriched motifs, edges indicate shared motif enrichment between modules. **D**) Enrichment of select miRNA targets within chrono-modules, based on miR Walk predictions. **E**) Network diagram of microRNA target enrichment within chrono-modules. Nodes indicate the number of enriched microRNAs per module, edges indicate shared microRNA enrichment between modules.

**Supplementary Figure 4.**
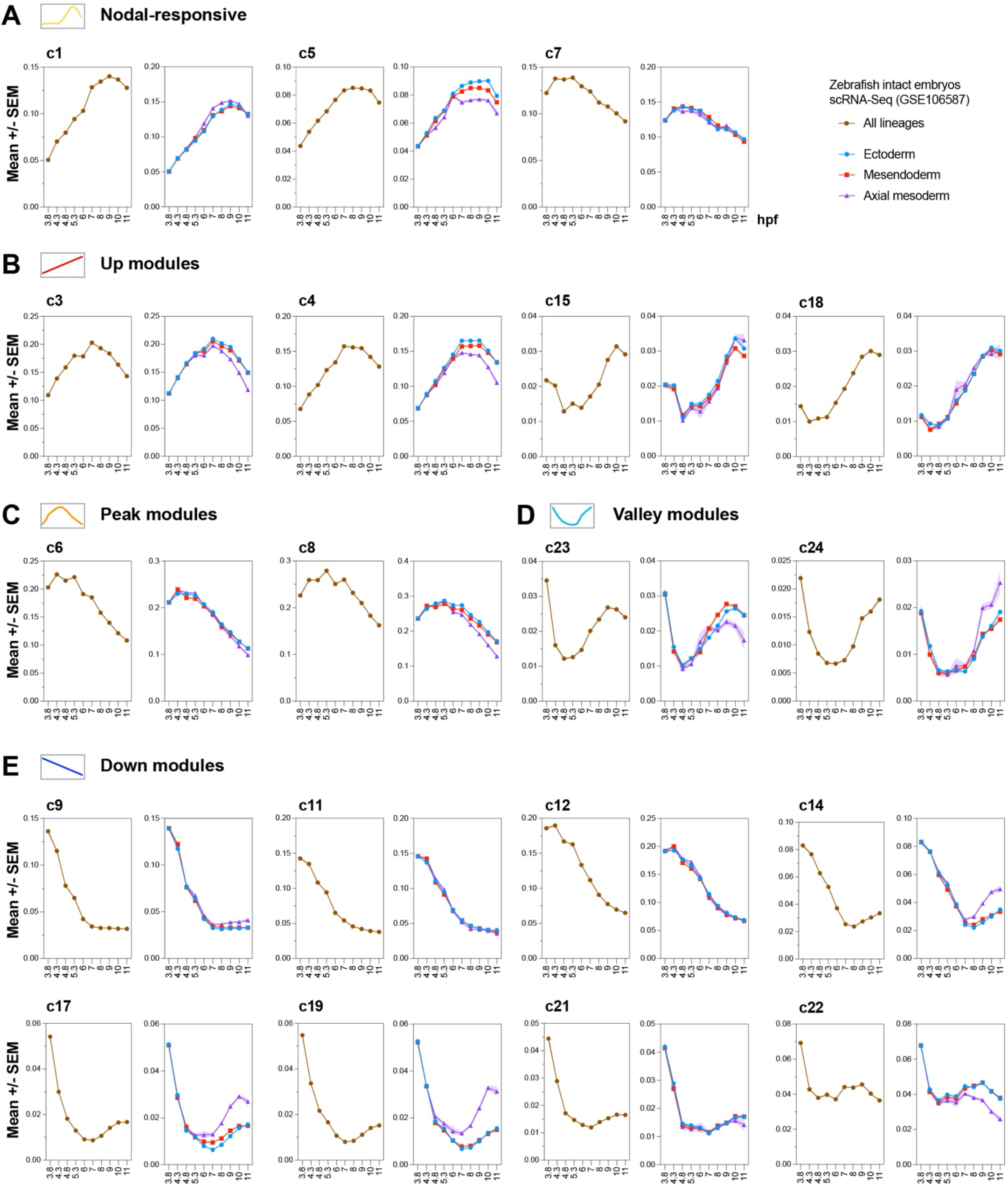
**Chrono-modules are concordant in pseudo-bulk and lineage-parsed single-cell data. A-E**) Summed z-score trajectories of the indicated chrono-modules for combined cell lineages (brown graphs) or for individual lineages (red= mesendoderm, purple= axial mesoderm, blue= ectoderm) from 3.8 to 11 hpf. Based on the single-cell zebrafish embryo atlas GSE106587 (*89*).

Many of these same TFBSs were also enriched within module 3, which exhibits an expression peak around gastrulation onset similar to that of module 12. Module 17 was moderately enriched for several Fox and ROR family TFBSs and module 23 for Yy1a/b binding sites. In some cases, expression of an implicated transcription factor within explants and intact embryos mirrored that of its putative target CCM. For example, the peaked expression of *etv5b* is similar to that of modules 3 and 12, and *yy1a* expression is a mirror image of the “valley” pattern of module 23 (**Fig. 4B**, compare with **Supp Fig. 1**), consistent with the (context dependent) role of Yy1 as a transcriptional repressor (*87, 88*). While a transcription factor’s transcript levels don’t necessarily reflect its activity, these striking patterns may support a role for specific transcription factors in regulation of specific CCMs. A network diagram of TFBS enrichment showed that chrono-modules with similar shapes (for example, the peaked shapes of modules 3, 8, and 12) were often enriched for the same putative transcriptional regulators (**Fig. 4C**). Notably, the Nodal-responsive module 16 shared no TFBSs with any CCMs.

MicroRNAs (miRNAs) also contribute to transcript levels via their ability to target mRNAs (especially maternally deposited mRNAs) for degradation (*80, 81*). We therefore examined our 24 modules for enrichment of predicted miRNA targeted transcripts using the miR Walk database (**Fig. 4D, Supp Table 3**). Most modules were moderately enriched for at least one miRNA, which we visualized using a co-targeting network (**Fig. 4E**). Strikingly, the module with the largest number of miRNAs predicted to target its constituent genes – module 13 – was the only module exhibiting monotonically decreasing expression. This likely reflects the large number of maternally expressed transcripts included in this module (**Supp Table 1**), which are known targets of miRNA-mediated degradation (*81*). The other modules putatively targeted by a moderate number of miRNAs were in the down, valley, and Nodal-responsive categories, while those with peaked or increasing expression were the predicted targets of very few miRNAs. This is consistent with the known function of miRNAs in generally reducing transcript levels and, together with TFBS enrichment, provides plausible mechanisms by which transcripts of a given expression module are temporally co-regulated.

### Chrono-constitutive modules are concordant in single cells isolated from zebrafish embryos

Findings from our explant system suggest that CCM gene expression varies strictly with developmental time rather than cell lineage. These genes exhibit similar expression patterns in intact embryos, but such bulk RNA-seq data cannot distinguish between different cell types and therefore cannot inform the cell type-specificity (or lack thereof) of CCMs in whole embryos. We therefore examined the temporal expression trajectories of genes belonging to all 24 modules in single cells isolated from zebrafish embryos across early embryonic stages (*89*). We first performed pseudo-bulk analysis of cells from ten stages between 3.8 and 11 hpf (**Fig. 5A**, top), in which we examined temporal expression trajectories of the 3936 chrono-module genes (out of 5000) present in this dataset. This analysis included 41 distinct cell types and 64 cell type/developmental stage pairs. For each cell type/stage pair with at least 100 cells, we conducted cell sub-sampling of 75 cells, then converted the data into pseudo-bulk representation. We then applied upper quartile normalization and z-score transformation and plotted summed z-scores for each CCM over developmental time. In many cases, pseudo-bulk CCMs closely resembled those in explants, in some cases better than the bulk whole embryo data, including modules 4, 15, 17, 19, 20, 21, and 22 (**Fig. 5, Supp Fig. 4**, brown graphs). In other cases, pseudo-bulk CCMs differed from those in explants or whole zebrafish embryos. For example, several peaked modules, including modules 6, 8, 9, 10, 11, 12, and 14, exhibited only the decreasing (right-hand) portion of the expression pattern in pseudo-bulk, similar to patterns observed in whole embryos (**Fig. 5, Supp Fig. 4**, brown graphs). These findings reflect a general concordance between temporal gene expression patterns in zebrafish embryo, explant, and single cell datasets.

**Figure 5.**
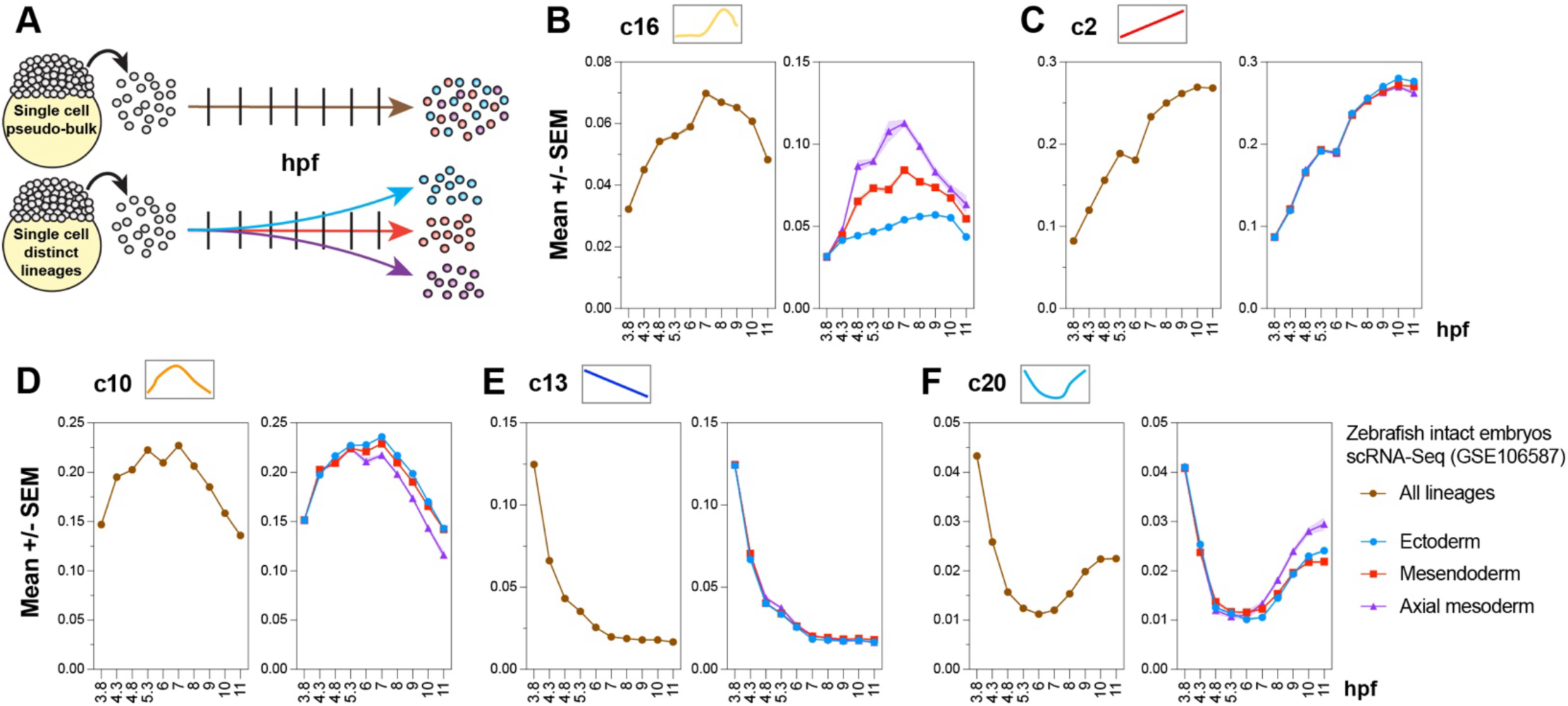
Chrono-modules are concordant in pseudo-bulk and lineage-parsed single-cell data. **A**) Schematic of single-cell RNA-seq analysis. We generated pseudo-replicates within each cell type and stage followed by pseudo-bulk representation of all lineages combined (brown) or in specific cell lineages (red, purple, blue). **B-F**) Summed z-score trajectories of the indicated chrono-modules for combined cell lineages (brown graphs) or for individual lineages (red= mesendoderm, purple= axial mesoderm, blue= ectoderm) from 3.8 to 11 hpf. Based on the single-cell zebrafish embryo atlas GSE106587 (*89*).

**Supplementary Figure 5.**
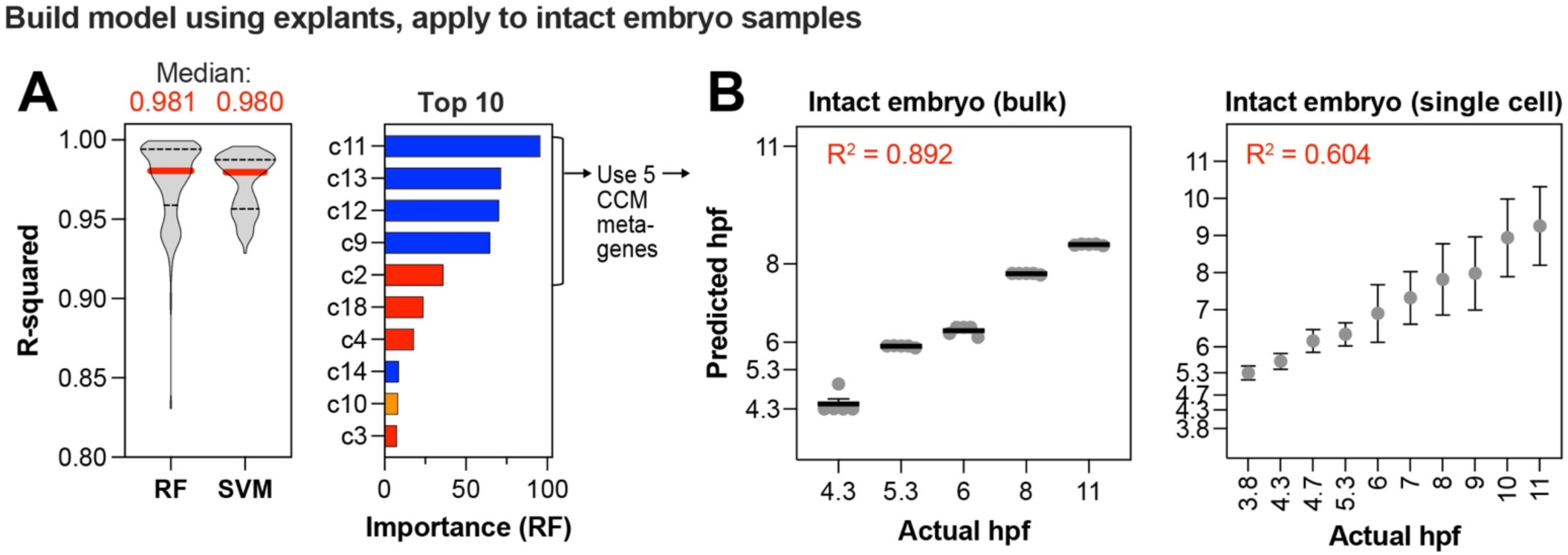
Machine learning predictors trained on bulk RNA-seq CCM UCell scores are less robust than single-cell-trained predictors. **A**) Machine learning predictors for developmental age (in hpf) were trained on UCell scores for all 20 CCMs in the explant bulk RNA-seq dataset. Performance of the Random Forest (RF) and Support Vector Machine (SVM) models in explants is shown as goodness-of-fit R^2^ over 100 cross-validation iterations (left). The top ten informative CCMs are shown for the RF model (right). **B**) A RF machine learning predictor trained on the top five CCM UCell scores was applied to bulk intact embryo RNA-seq (left) and single-cell RNA-seq (right) datasets. Scatterplots show predicted vs. actual hpf and goodness of fit (R^2^) of the predictions.

To test our hypothesis that CCM genes identified in explants exhibit consistent temporal expression trajectories between distinct tissue types within whole embryos, we further divided cells from this single-cell dataset into each of three major lineages: ectoderm, mesendoderm, and axial mesoderm (**Fig. 5A**, bottom). These lineages exhibit the clearest distinction in reconstructed developmental trajectories, and split from one another at the first major branchpoint among embryonic cells (*89*). We applied our analytical strategy to each lineage, including sub-sampling by cell type and stage, pseudo-bulk transformation, upper quartile normalization, and plotting summed z-scores for each CCM. Examining these plots, we observed that expression trajectories were strikingly consistent between all three cell lineages (**Fig. 5C-F**, **Supp Fig. 4**, red/blue/purple graphs). The clearest exception is the Nodal-responsive module 16 (**Fig. 5B**), in which expression trajectories varied greatly between the three cell linages. Because the presence and levels of Nodal signaling are largely responsible for specifying these different lineages (*90, 91*), this divergence is expected and serves as validation of our approach. Surprisingly, however, expression trajectories in the Nodal-responsive modules 1 and 7 did not differ substantially between lineages (**Supp Fig. 4A**). For most CCM modules, expression trajectories were nearly identical between each of the three lineages and all cells in the pseudo-bulk data (**Fig. 5**, **Supp Fig. 4)**, but a few modules did exhibit small inter-lineage differences. Interestingly, among such modules, the ectoderm and mesendoderm tended to be similar while the axial mesoderm diverged at later time points (**Fig. 5D, F**). Overall, these results are consistent with our hypothesis that CCM gene expression varies strictly with developmental time in a cell type-agnostic manner, highlighting a potential role for these genes in embryo-wide timekeeping.

Although our CCM trajectory analysis using pseudo-bulk data was consistent with the results obtained in explants and intact embryos, it still lacked true single-cell resolution. A known limitation of single-cell RNA-seq data is the stochastic nature of the genes detected, with considerable gene drop-out and variation in genes detected cell by cell (*92*). Because such data would not be amenable to detection of CCMs by summed z-scores, we sought to instead identify CCM gene signatures for each cell. We therefore applied UCell (*93*), which uses a non-parametric approach to summarize the genes detected in each individual cell into a single gene signature score based on the Mann-Whitney U statistic. To enable direct comparisons between bulk and single-cell RNA-seq data, we first filtered each dataset by the 5000 most variable genes detected in our explants. We then applied the UCell score to the full single-cell dataset, composed of 34,016 cells, as well as to each of the 63 explant and 25 intact embryo bulk RNA-seq samples. We plotted the chrono-module scores over time (**Fig. 6A-C**) and observed a robust concordance not only of UCell trajectories between datasets, but also between UCell and summed z-score trajectories within each dataset. Strikingly, the single-cell dataset refined the trajectories and identified deviations between certain stages not observed in other datasets, indicating single-cell properties that are missed by bulk RNA-seq analysis.

**Figure 6.**
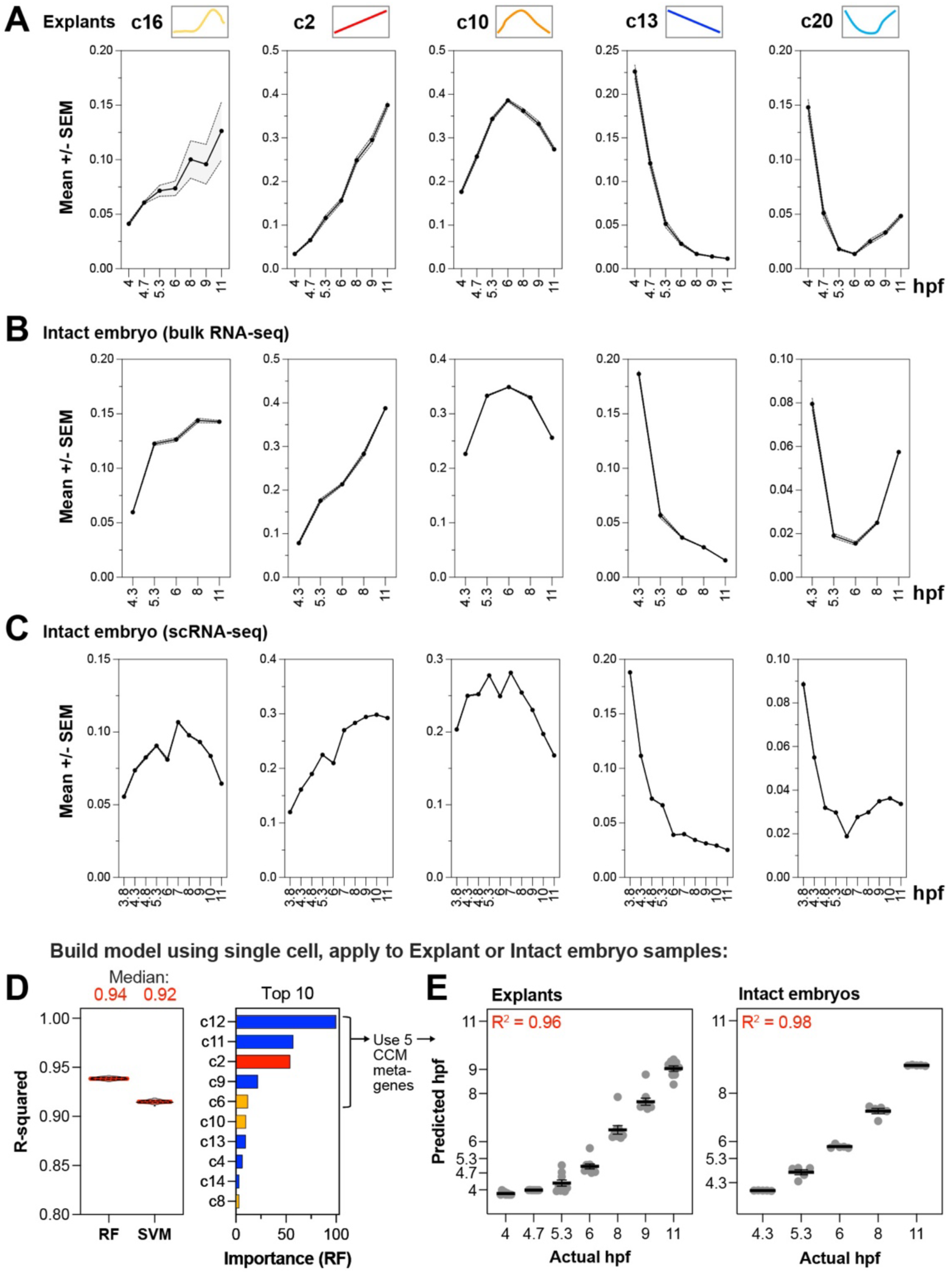
UCell scores enable comparisons of chrono-module trajectories within individual cells and accurate prediction of developmental age. A-C) A UCell score was computed for every chrono-module at each timepoint within bulk explant RNA-seq (A), bulk intact embryo RNA-seq (B), and each cell of the single-cell zebrafish embryo dataset (C). The distribution of UCell scores across developmental time is shown for select chrono-modules. **D**) Machine learning predictors for developmental age (in hpf) were trained on UCell scores for all 20 CCMs in the 34,016-cell scRNA-seq dataset. Performance of the Random Forest (RF) and Support Vector Machine (SVM) models in single cells is shown as goodness-of-fit R^2^ over 100 cross-validation iterations (left). The top ten informative CCMs are shown for the RF model (right). **E**) A RF machine learning predictor trained on the top five CCM UCell scores was applied to bulk explant RNA-seq (left) and bulk intact embryo RNA-seq (right) datasets. Scatterplots show predicted vs. actual hpf and goodness of fit (R^2^) of the predictions.

**Supplementary Figure 6.**
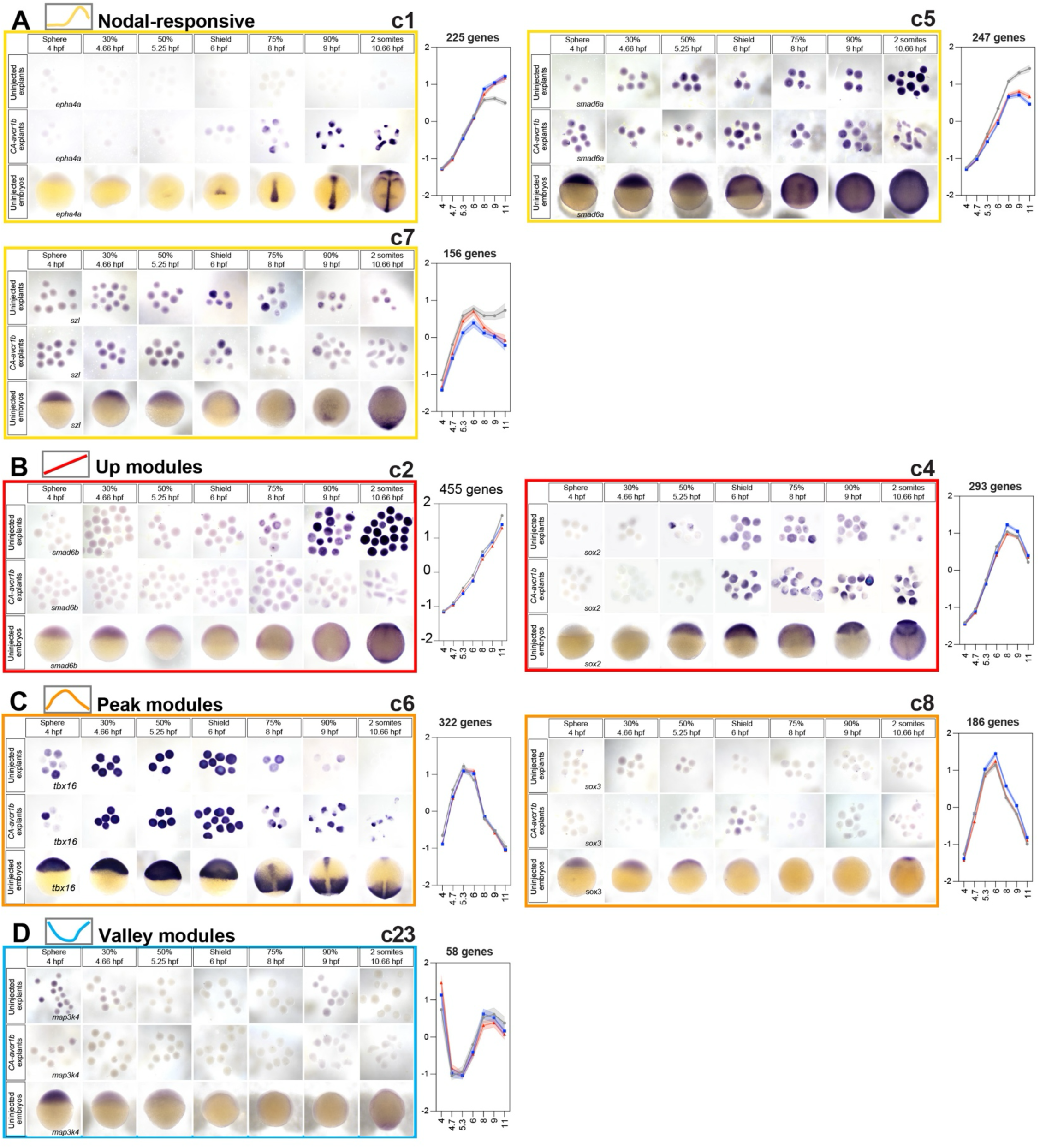
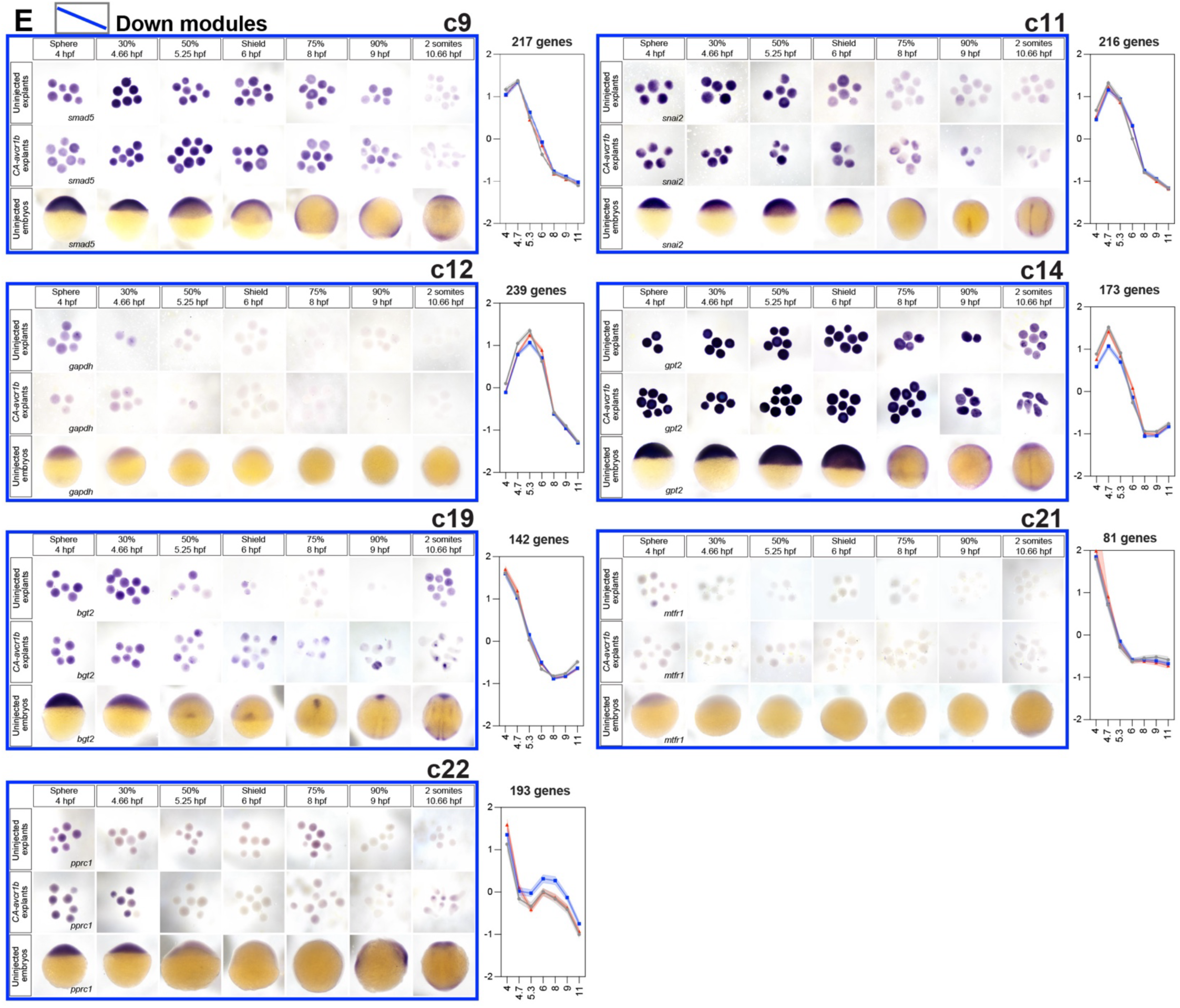
***In vivo* expression patterns largely recapitulate chrono-module expression trajectories. A-E**) Whole mount *in situ* hybridizations for a representative gene (indicated in left-most panels) of each chrono-module in uninjected explants (top), *CA-acvr1b*\* explants (middle) and intact zebrafish embryos (bottom) at the stages indicated. Each staining pattern is shown alongside the summed Z-score trajectories for the module to which the gene belongs.

### Expression of chrono-constitutive genes accurately predicts stage in zebrafish explants and embryos

Our analysis has revealed a set of temporally dynamic genes, clustered into 20 distinct modules, whose expression trajectories are largely consistent across teleost species, cell types, and experimental perturbations (like explantation and Nodal activation). We hypothesize that these genes form the basis of a transcriptional timer, steadily progressing with developmental stage across all embryonic cell lineages. If these genes in fact function this way, we further hypothesize that expression levels of CCM genes would provide sufficient timing information to stage embryos and embryonic cells based on transcriptomes alone.

To this end, we applied a previously published machine learning pipeline (*94-96*) - including Random Forest (RF) and Support Vector Machine (SVM) approaches - to the UCell scores of the single-cell RNA-seq data. For each of 100 iterations, we divided the data into an 80% training set from which we built the models, and applied them to a 20% test set for which we computed the goodness-of-fit metric, R^2^. These CCM machine learning models predicted hpf of individual embryonic cells with a median R^2^ of 0.94 and 0.92 for RF and SVM, respectively, indicating very strong modeling performance (**Fig. 6D**). Using the Interpretable Machine Learning (IML) approach, we determined that the CCMs most informative for age prediction by the RF model were 12, 11, 9 (all down modules), 2 (an up module), and 6 (a peak module) (**Fig. 6D**). We next asked whether these five CCMs are sufficient to predict embryonic age in other datasets. Remarkably, using an RF model built using only the scRNA-seq UCell scores of the five most informative CCMs, we predicted hpf of the 63 explant bulk RNA-seq samples with R^2^=0.96 and of the 25 intact embryo bulk RNA-seq samples with R^2^=0.98 (**Fig. 6E**). These results demonstrate that just a few CCM metagenes are sufficient to accurately predict the embryonic age of zebrafish embryos and even individual embryonic cells, highlighting their ability to confer developmental timing information.

We also trained a machine learning model on CCM UCell scores from our explant dataset, which achieved strong performance (R^2^=0.981 for RF and 0.980 for SVM) when tested on explants (**Supp Fig. 5A**). Notably, the most informative CCMs for this model were similar to those for the single-cell model, with “down” modules being overrepresented (**Supp Fig. 5A**). This explant-trained model predicted the age of intact embryos (bulk RNA-seq data) reasonably well with R^2^=0.892, but its performance predicting the age of the 34,016 individual cells was moderate, with R^2^=0.604 (**Supp Fig. 5B**). This indicates that although machine learning models trained on CCMs from bulk RNA-seq datasets are informative for other bulk RNA-seq datasets, single-cell prediction models are more accurate and widely applicable, presumably because they are trained on larger datasets with higher intercellular variability.

These models demonstrate that transcriptomic data alone can be used to predict the actual developmental age of zebrafish embryos, explants, and even individual cells, and that sufficient staging information can be conferred by as few as five metagenes. Furthermore, such staging uses only chrono-constitutive genes whose expression is consistent across cell lineages, i.e. it requires no markers of cell type-specific differentiation, and is accordingly robust in individual cells regardless of lineage. Together, our findings are consistent with a core set of timekeeping genes whose expression reliably marks the passage of developmental time across all cell types within an embryo.

### In vivo validation of CCM gene expression dynamics

Our work identified gene modules that, according to RNA-seq data, exhibit distinct temporal expression trajectories. To validate these patterns within zebrafish embryos and explants, we performed colorimetric wholemount *in situ* hybridization (WISH) for one representative gene from most chrono-modules within Nodal-activated explants, uninjected explants, and intact embryos at the seven developmental stages analyzed by RNA-seq (**Fig. 7**). Among these select examples, we found that many recapitulate both the temporal expression trajectories and similarities or differences between conditions shown in the RNA-seq data. For example, representative genes from all four Nodal-responsive modules were visibly different between Nodal-activated and uninjected explants, with those from modules 16 and 1 being higher in Nodal-activated and those from modules 5 and 7 being higher in uninjected explants, as predicted (**Fig. 7A, Supp Fig. 6A**). Genes within CCMs also generally exhibited the up, down, peak, and valley expression trends by which they were clustered (**Fig. 7B-E, Supp Fig. 6B-E**).

**Figure 7.**
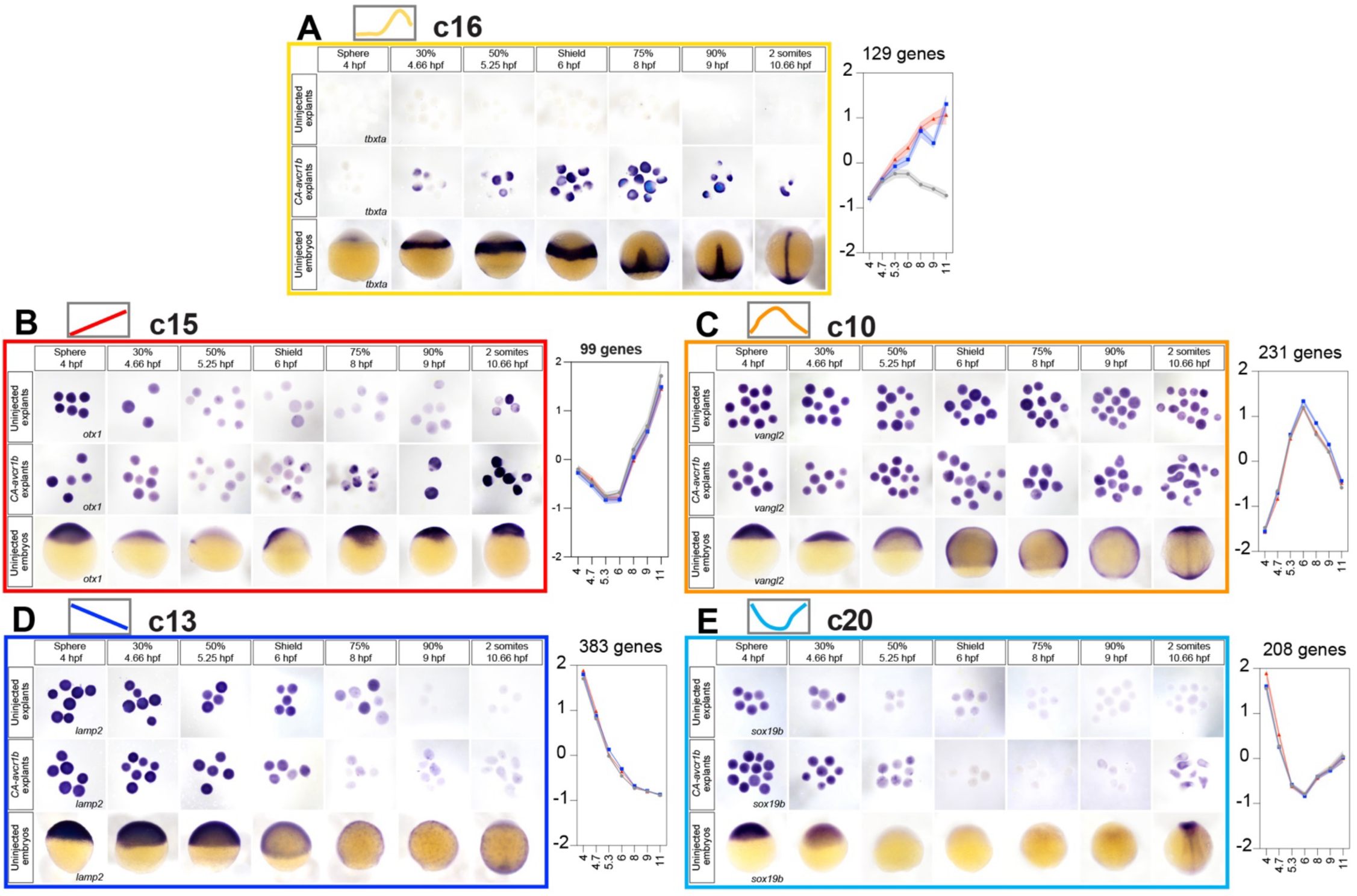
*In vivo* expression patterns largely recapitulate chrono-module expression trajectories. **A-E**) Whole mount *in situ* hybridizations for a representative gene (indicated in left-most panels) of each chrono-module in uninjected explants (top), *CA-acvr1b*\* explants (middle) and intact zebrafish embryos (bottom) at the stages indicated. Each staining pattern is shown alongside the summed Z-score trajectories for the module to which the gene belongs.

As mentioned above, we did see examples of CCM genes whose temporal expression trajectories were generally similar between conditions but with different absolute expression levels. This was evident for *sox2* and *smad6b,* whose expression increased in all conditions but was more strongly expressed in Nodal-induced or uninjected explants, respectively (**Supp Fig. 6B**). It was also seen for *sox19b* and *tbx16*, whose increased expression on the right-hand side of the “valley” or “peak”, respectively, was higher in Nodal-activated explants and intact embryos than in uninjected explants (**Fig. 7E, Supp Fig. 6C**). We also observed that not all CCM genes exhibited ubiquitous expression throughout the developmental time-course, as we might expect for genes with cell type-agnostic expression. Indeed, many of the representative genes we selected for WISH were expressed ubiquitously at early stages but later resolved into tissue-specific expression patterns (see for example *tbx16*, *snai2*, *gpt2*, and *bgt2* (**Supp Fig. 6**)). The reason for this is likely two-fold. First, because uninjected explants contain two cell types (non-neural ectoderm and enveloping layer), some genes that we identified as chrono-constitutive may be specific to one of these two cell types. Our cell lineage-specific analysis of single-cell data (**Fig. 5**) suggests this is not the case for the majority of CCM genes, however.

Which brings us to the second reason: genes with cell type-specific expression are likely overrepresented among our *in vivo* validated examples because they are more likely to have robust WISH probes available through the zebrafish international resource center (ZIRC). Even among examples with tissue-specific expression *in vivo*, however, their expression levels in explants of both conditions largely recapitulated those expected from RNA-seq and thus, validated temporal expression patterns of chrono-module genes.

## Discussion

How embryos keep time is a fundamental but poorly understood question in developmental biology. Here, we have begun to address this question by identifying transcriptomic changes that are chrono-constitutive, that is, that vary with developmental time in a cell type-agnostic way. We identified such genes by 1) comparing longitudinal gene expression in zebrafish explants containing limited cell types to those containing a full complement of germ layer derivatives, and 2) comparing expression levels over time between distinct cell lineages from an embryonic zebrafish single-cell RNA-seq dataset. Both approaches revealed that, contrary to our initial expectations, most dynamic gene expression changes across early zebrafish development are stage - rather than tissue - dependent. This is consistent with the recent finding that levels and dynamics of individual transcripts are highly similar between individual cells of different lineages (*54*). Further, clusters of temporal expression trajectories, which we term chrono-modules, match remarkably well between *ex vivo* (explant) and *in vivo* (embryo) datasets. Together, these analyses have uncovered a set of strictly “timekeeping” genes whose expression reflects developmental time independent of specification or differentiation of any particular cell type.

### How much of embryonic gene expression is truly chrono-constitutive?

Our analyses support the conclusion that dynamic expression of most genes varies with developmental time rather than cell type during early zebrafish development. However, lineage-specific analysis and validation of select expression patterns by WISH revealed that some genes we categorized as chrono-constitutive in fact exhibited tissue-specific expression during at least one stage examined (**Supp Fig. 4**, **Supp Fig. 6**). Further, although our single-cell analysis showed that most CCMs had nearly identical temporal expression trajectories across distinct cell lineages (**Fig. 5**), expression within the axial mesoderm diverged slightly from the other lineages in some modules (**Supp Fig. 4**). This may reflect, at least in part, the smaller number of axial mesoderm cells (∼2,000) than of ectoderm and mesendoderm (∼13,000 and 10,000, respectively) used in our analysis. Notably, if CCM expression diverged between cell lineages, it did so at later time points (**Supp Fig. 4**) when tissue-specific expression of some genes became apparent in whole embryos by WISH (**Supp Fig. 6**).

This suggests that explants may not fully recapitulate the tissue-specific expression patterns of all genes in intact embryos, especially during late gastrulation. We speculate that such tissue-specific genes were also overrepresented among those we validated by WISH because those genes with robust probes available from public resource centers tend to be those with “interesting” (i.e. tissue-specific) expression patterns. Together, this highlights a limit of our methods, which assumed that gene expression in uninjected explants was cell-type agnostic, leading to a slight overestimation of the number of chrono-constitutive genes. Additional gene-by-gene screening for consistent expression across all cell lineages within single cells would enable us to filter out these not strictly chrono-constitutive genes.

### The utility of chrono-constitutive gene expression for embryonic staging

Despite the presence of some not strictly chrono-constitutive genes, we demonstrated the utility of our CCMs for predicting embryonic time by generating machine learning models trained thereon. Indeed, the modules identified as most important to the ability of our machine learning models to predict stage were 2, 11, and 12 (**Fig. 6**), three CCMs with essentially no inter-lineage differences in our single-cell analysis (**Fig. 5A, Supp Fig. 4E**). Several previous bioinformatic approaches accurately ordered developmental gene expression profiles from earliest to latest (*37, 38, 47, 48*). However, this reflects pseudo-time rather than real chronological time, age, or stage. The tool RAPToR, by contrast, performs real age (stage) prediction by comparing transcriptomes from discrete stages to reference datasets, and can even accurately stage embryos of one species using reference data from another (*49*). However, muscle differentiation signatures provided the majority of staging information (*49*), limiting its utility to intact embryos but not cells from other lineages. In contrast, single-cell sequencing enables identification of timekeeping genes common to all cell types. Several single-cell analysis tools used transcriptomic aging/staging profiles to assign individual cells to broad stage categories (*50*), improve developmental lineage trees (*51*), determine the “differentiation stage” of individual cells (*52*), or compute “transcriptional ages” for each tissue type within an embryo (*53*). Many of these single-cell staging approaches estimate “phylogenetic”, “differentiation”, or “pseudo” time, whereas our CCM-based model predicts absolute chronological age/ stage. Further, our model trained only on CCMs in single cells predicted the stage of explants and intact zebrafish embryos, demonstrating the sufficiency of chrono-constitutive genes to infer developmental time.

This raises the fascinating question of whether the embryo uses CCMs to keep its own developmental time, or whether such temporal expression patterns are simply a byproduct of developmental progression without an instructive role therein. It is similarly unclear whether CCMs govern developmental progression, respond to it, or both. Addressing these questions in the future will require methods to modify CCM gene expression or developmental progression to examine how each responds to changes in the other, and how each affects the milestones of embryonic development.

### Regulation of timekeeping gene expression

Another important open question is how CCM gene expression is regulated. This study identified candidate transcriptional and post-transcriptional regulators for several modules, but future experimental studies will be needed to examine the role of these candidates in CCM expression trajectories. It is also unclear how all CCMs are coordinated with one another and with the progression of developmental time. For example, are all CCMs under the control of some “master” timer? Do some govern the expression of others through a series of feedback interactions? Or is the expression of every (or any) CCM independent of all others? Distinguishing between these possibilities will require experimental strategies to disrupt individual CCMs one at a time and quantify their effect of trajectories of others. If CCMs are indeed at least partially independent, it would raise the possibility that discrete developmental events (for example, loss of pluripotency or onset of morphogenesis) are under the control of distinct CCMs. Such relationships may be revealed by careful examination of developmental milestones upon disruption of a given CCM in the future.

### Evolutionary conservation of timekeeping genes

Excitingly, our analysis revealed that CCMs are largely conserved in another teleost fish, the Japanese medaka. This suggests that common transcriptional programs may mark (and perhaps govern) developmental time across a variety of species. Indeed, even distantly related mice and *Drosophila* were found to share temporal expression patterns of some gene modules, including those involved in both cell type-specific (muscle differentiation) and cell type-agnostic (cell cycle, RNA metabolism) processes (*25, 65*). Further comparisons between additional species will reveal how well conserved chrono-constitutive gene expression is, and may provide further insights into the roles of such genes in the timing of specific developmental events and/or developmental progression in general.

## Methods

### Zebrafish

Adult zebrafish were maintained through established protocols (*97*) in compliance with the Baylor College of Medicine Institutional Animal Care and Use Committee. Embryos were obtained through natural mating and staging was based on established morphology (*98*). Studies were conducted using embryos of the AB wild type strain. Fish were crossed from their home tank at random and embryos were chosen for injection and inclusion in experiments at random. Embryos were collected and fixed in 4% PFA upon reaching each of our seven desired timepoints.

### Blastoderm explants

Blastoderm explantation was performed according to (*61*). Embryos were dechorionated with pronase (1 mL of 20 mg/mL stock in 15 mL 3X Danieau’s) at 128 cell stage and washed with egg water and 0.3X Danieau’s solution. Explants were cut using Dumont #55 forceps (Fisher Science #NC9791564) on an agarose coated 60 mm X 15 mm plate filled with 3X Danieau’s solution. After cutting, explants were allowed to heal in the 3X Danieau’s solution for 5 minutes before being placed in agarose coated 6-well plates filled with explant media (DMEM/F12 Thermo #11330032, 3% Newborn calf serum, 1:200 penicillin-streptomycin). Explants were collected and fixed in 4% PFA when stage matched intact embryos reached each of our seven desired timepoints.

### Whole mount in situ hybridization

Antisense riboprobes were transcribed using NEB T7 or T3 RNA polymerase (NEB #M0251s and Fisher #501047499) and labeled with digoxygenin (DIG) NTPs (Sigma/Millipore #11277073910). Whole mount *in situ* hybridization (WISH) was performed according to (*99*) with minor modifications. Embryos and explants were fixed as described above, washed in PBS + 0.1% Tween-20 (PBT), gradually dehydrated, and stored in methanol at -20C. Immediately prior to staining, embryos were rehydrated into PBT and hybridized overnight with antisense probes within the wells of a 24-well plate. Samples were gradually washed into SSC buffer and then into PBT before overnight incubation with an anti-DIG primary antibody at 1:5000 (Roche #11093274910). Samples were washed in PBT and then staining buffer before developing in BM Purple staining solution (Roche #11442074001), then washed and stored in stop buffer (10 mM EDTA in PBT) until imaging. WISH images were taken with a Nikon Fi3 color camera on a Nikon SMZ745T stereoscope.

### RNA-seq processing

Bulk RNA-seq data from datasets GSE246158 and PRJEB12982 a were mapped using STAR (*100*) onto the zebrafish danRer11 genome assembly. Gene expression was quantified using featureCounts (*101*). Transcripts were converted to counts per million (CPM) then normalized using upper quartile normalization using the package EdgeR (*102*).

### Discovery of chrono-modules

The top 5000 most temporally variable genes were detected for the 63 explant bulk RNA-seq samples, then genes were z-score normalized. Consensus clustering was applied to the normalized data using the R package ConsensusClusteringPlus (*103*), generating 2 to 24 clusters. The clustering hierarchy was plotted using the clustree package (*104*). Module scores were computed for each sample and each chrono-module by summing the z-scores across all genes within the module. Module trajectories were plotted across the developmental stages profiled.

### Processing of published intact embryo RNA-seq data

Single-cell data read counts and metadata were downloaded for GSE106587 (*89*). Data were processed and normalized using Scanpy (*105*). For pseudo-bulk analysis, cells were first grouped by cell type and developmental stage. Cell type/stage pairs with at least 100 samples were subsampled 5 times, each time selecting randomly 75 cells; data were further converted to pseudo-bulk representation using the Python package decoupler (*106*). This process was applied for combined cells of all lineages, and separately for cells from the distinct lineages ectoderm, mesendoderm, and axial mesoderm. Data were further normalized using upper quartile normalization using the package EdgeR (*102*). Genes were then z-score transformed, and chrono-module scores were computed using summed z-score for each pseudo-bulk pseudo-replicate.

### Single-cell level hpf prediction

Bulk RNA-seq data from explants (GSE246158) and intact embryos (PRJEB12982) were processed as single cells using Scanpy (*105*). For all datasets, data were converted from AnnData/h5ad format to Seurat/rds format using zellkonverter (*107*). UCell scores were computed for each CCM using the UCell R package (*93*). We used the 34,016 single cells as discovery data; we used a cross-validation strategy, where for each iteration data were split into 80% training and 20% testing subsets. Random Forest (RF) and Support Vector Machine (SVM) methods were used to derive predictors, which were then applied to the 20% testing dataset. The goodness-of-fit R^2^ method was used to assess model performance in the testing set. This approach was used for 100 cross-validation iterations, then the median R^2^ was computed. Feature importance was determined for each CCM using the Interpretable Machine Learning (IML) R package (*108*) for each cross-validation iteration, and median values for each CCM were computed across all the cross-validation iterations. Predictors were then applied to the explants and intact embryo RNA-seq data using the UCell scored representation.

## Data access

The RNA-seq data for the zebrafish explant time-course was previously published (*59*) and has been deposited to GEO, accession GSE246158.

## Research funding

This work was supported by R01 HD104784 to MKW and MH134392 to CC. RK, SLG, and DV were partially supported by CPRIT RP210227 and RP200504, NIH/NCI P30 shared resource grant CA125123, NIH/NIEHS P42 ES027725 and P30 ES030285.

## Acknowledgements

We thank the BCM Center for Comparative Medicine for the care of our fish and Dr. Lila Solnica-Krezel for sharing WISH probes. Thanks also to all members of the Williams and Coarfa labs and Drs. Shelby Blythe, Nathan Lord, and Jeff Farrell for helpful discussions.

